# A 14-Color Blue-Violet Laser Restricted Full Spectrum Flow Cytometry Panel for Comprehensive Immunophenotyping of Pulmonary Inflammation in Mouse Bronchoalveolar Lavage Fluid

**DOI:** 10.64898/2026.01.23.701343

**Authors:** Arlind B. Mara, Jeremy M. Miller, R. Grace Ozyck, Morgan L. Hunte, Edan R. Tulman, Steven M. Szczepanek, Steven J. Geary

## Abstract

Here we describe a 16-parameter, 14-color surface staining panel optimized for murine bronchoalveolar lavage cells that enables reproducible identification of major innate and adaptive immune populations relevant to pulmonary infections or other inflammatory conditions of the airways. The panel enables confident identification of neutrophils, eosinophils, B cells, T cells and subtypes, NK cells, and distinguishes between tissue resident and monocyte-derived macrophages populations. The panel was carefully designed for BAL samples that vary in cell number, are rich in debris, and often autofluorescent. Antibody concentrations are optimized to provide reproducible results regardless of sample-variable cell numbers allowing for the preparation of a single antibody cocktail master mix and rapid sample staining time, thereby cutting down on sample preparation and optimizing cell viability of analyzed samples. The panel facilitates robust cross-sectional and longitudinal comparison of airway inflammation across different airway inflammatory conditions, infections by different respiratory pathogens, impact of vaccination or therapeutics on the inflammatory landscape, and more. It facilitates hypothesis generation by revealing recruitment kinetics and remodeling of myeloid compartments, supports downstream sorting for transcriptomic or functional assays, and provides a standardized baseline for labs to adopt or extend for activation or intracellular cytokine analyses. We have successfully utilized this panel to identify differential host responses to different respiratory Mycoplasma pathogens as well as to longitudinally track the progression of inflammatory response to Mycoplasma pneumoniae over a 21-day time course study. This panel provides an economic immunophenotyping option by utilizing only 14 markers and a two laser (Blue and Violet) full spectrum cytometer to provide comprehensive immunophenotyping power of both myeloid and lymphoid cells. Furthered by lacking the requirement for advanced unmixing for sample analysis, the panel can be easily adopted by the community, enabling comparative meta-analyses of host responses across murine respiratory infection models.

## Background

Immunophenotyping by flow cytometry yields high-throughput, single-cell resolution of the inflammatory landscape in complex tissues and fluids, making it an essential tool for dissecting host-pathogen interactions in the lung. Bronchoalveolar lavage fluid (BALF) samples provide a proximal readout of the airway and alveolar immune compartments and thus reflect the combined outcomes of pathogen sensing, innate effector recruitment, adaptive activation, and tissue repair processes^1-4^. Quantifying which cell types are recruited and how their relative abundances change over time in conjunction with how these changes correlate with pathogen burden, cytokine milieu, and histopathology enables us to infer mechanistic drivers of clearance, persistence and immunopathology. Immunophenotyping of BALF inflammation is particularly valuable for comparative pathogen studies as different pathogens and strains may provoke distinctive immune landscapes that underlie differences in pathogenicity, virulence, tissue damage, and vaccine responsiveness. For example, early neutrophil predominance often indicates rapid innate recruitment and potential tissue-damaging inflammation, whereas eosinophil or monocyte-derived macrophage signatures often point to alternative innate pathways associated with tissue reparative phenotypes or type-2 skewing^5-8^. Furthermore, time course profiling of the inflammatory landscape can reveal recruitment and resolution kinetics that implicate chemokine networks and cellular differentiation pathways which can guide targeted mechanistic experiments such as cytokine neutralization, cell depletion, or single-cell transcriptomic experiments for more in depth immunophenotyping.

BALF samples present technical challenges that make traditional panel design difficult and limit the ability for downstream sample analysis using dimensionality reduction approaches. One such issue is that cell yields are often low and variable, slowing down sample preparation time by requiring adjustments of antibody concentrations to cell count to reduce artifacts of staining for downstream analysis^3,4^. Furthermore, debris in BALF samples can obscure small populations, and alveolar macrophages exhibit high autofluorescence that can make cell identification difficult^3-4,9-10^. By utilizing a full-spectrum approach which takes advantage of autofluorescence extraction and optimizing antibody concentrations and staining protocols to provide robust signal when staining samples with variable cell numbers, our panel provides an economical, accessible, and easy to use alternative to the broader larger panels^11^ that enables reliable and robust identification of major mouse leukocytes involved in inflammatory phenomena.

## Purpose and Applications

Our primary reason for developing this panel was to provide a single, validated 14-color surface panel for routine, comparative immunophenotyping of murine BALF cells in models of Mycoplasma pulmonary infections. Indeed, we have successfully utilized this panel to identify differential host responses to different respiratory Mycoplasma pathogens as well as to longitudinally track the progression of inflammatory response to *Mycoplasma pneumoniae* over a 21-day time course. We see this panel as being of utility to researchers looking to conduct cross-sectional or longitudinal studies that compare inflammatory phenotypes of the lung environment due to different pathogens, pathogen strains, host genotypes, or to assess the effect of certain therapeutics or prophylactics on host response to infection. Examples of applications can include profiling acute versus resolving inflammation during infection, screening vaccine or therapeutic effects on cell recruitment kinetics, as well as hypothesis generation for targeted follow up studies such as cell sorting for transcriptomics, intracellular cytokine staining, cell depletion experiments, cytokine neutralization experiments etc. Beyond infectious models, the panel can be broadly useful for assessing allergic airway disease, COPD, effect of respiratory exposures to environmental or occupational toxins, sterile lung injury and chronic remodeling of fibrotic processes as it resolves the key leukocytes central to these conditions^5-8,12-16^ . When correlated with pathogen burden, cytokine profiles, histopathology, or lung function, immunophenotyping by this panel can accelerate mechanistic and translational work on pulmonary conditions that burden human and animal health.

## Overall Experimental Design and BALF Collection

Mice were anesthetized via vaporized isoflurane then intranasally inoculated with 1x10^8 CFU of appropriate *Mycoplasma* inoculum in 50uL of growth medium, or 50uL of sterile growth medium to serve as controls. At the appropriate time point post infection (4 days for comparative study, 3, 6, 9, 12, 15, 18, and 21 for time-course study), mice were anesthetized as above and humanely euthanized via cervical dislocation in accordance with our approved IACUC protocol #A24-024. Aseptically inside a class 2 biosafety cabinet the trachea was exposed by dissecting away tissue and the BALF was collected by using 1.5mL of sterile PBS loaded onto a 3mL syringe equipped with an half-inch 18G needle. The needle was inserted into the lumen of the trachea until the bevel was fully covered, the needle affixed on the trachea using forceps, and the lungs were lavaged by gently aspirating 1mL of the 1.5mL PBS back and forth 3 times. The fluid was collected onto a sterile, capped FACS tube which was then placed until all experimental samples were collected.

## Sample Processing and Staining

Following sample collection, the cells of the BALF fluid were pelleted via centrifugation at 500 rcf (xg) for 10 minutes at 4C. The supernatant was aspirated and collected for archival purposes, and the cells were resuspended in 200uL of homemade blocking FACS buffer containing the following: *(1x Hank’s Balanced Salt Solution -phenol red free (HBSS), 10% Rabbit Serum, 0*.*9% Sodium azide, 5ug/mL of purified anti-mouse CD16/32 antibody (clone 93))*.Following resuspension, the cells were incubated on ice and in the dark for 5 minutes to block. The Antibody Staining MasterMix (MM) four our panel (**Table 1**) was prepared in blocking homemade FACS buffer to the concentrations listed in **Table 2** and **3** below, based on concentrations determined through serial dilutions displayed on **Figure 1**. Following 5 minutes of blocking, 50uL of MM was added to each sample, and the samples were gently mixed before placing back on ice, in the dark to stain for 15 minutes. Following staining, 3mLs of FACS buffer (*as blocking FACS buffer above but without added anti-mouse CD16/32 blocking antibody*) was added to each tube to wash cells. Cells were pelleted by centrifugation as above and resuspended in 2oouL of FACS buffer per sample and stored on ice in the dark. 100uL of 1x Sytox Blue Live-Dead dye was added to the sample 30 seconds prior to acquisition to stain dead cells, and samples were acquired at a flow rate between 500-4000 events per second.

**Table 1.**
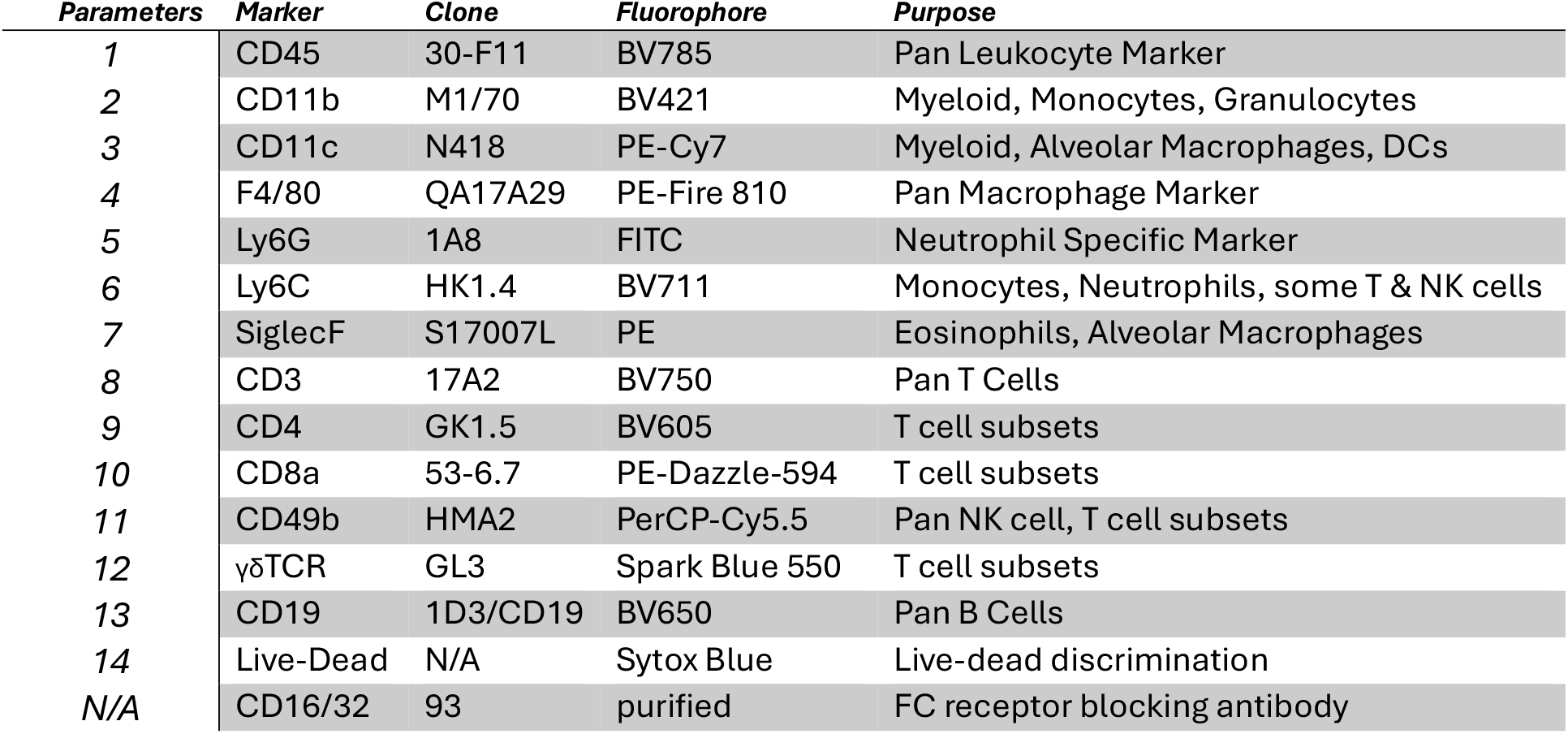
Reagents used for optimized multicolor panel and leukocyte identification.

| *Parameters* | *Marker* | *Clone* | *Fluorophore* | *Purpose* |
| --- | --- | --- | --- | --- |
| 1 | CD45 | 30-F11 | BV785 | Pan Leukocyte Marker |
| 2 | CD11b | M1/70 | BV421 | Myeloid, Monocytes, Granulocytes |
| 3 | CD11c | N418 | PE-Cy7 | Myeloid, Alveolar Macrophages, DCs |
| 4 | F4/80 | QA17A29 | PE-Fire 810 | Pan Macrophage Marker |
| 5 | Ly6G | 1A8 | FITC | Neutrophil Specific Marker |
| 6 | Ly6C | HK1.4 | BV711 | Monocytes, Neutrophils, some T & NK cells |
| 7 | SiglecF | S17007L | PE | Eosinophils, Alveolar Macrophages |
| 8 | CD3 | 17A2 | BV750 | Pan T Cells |
| 9 | CD4 | GK1.5 | BV605 | T cell subsets |
| 10 | CD8a | 53-6.7 | PE-Dazzle-594 | T cell subsets |
| 11 | CD49b | HMA2 | PerCP-Cy5.5 | Pan NK cell, T cell subsets |
| 12 | $\gamma\delta$TCR | GL3 | Spark Blue 550 | T cell subsets |
| 13 | CD19 | 1D3/CD19 | BV650 | Pan B Cells |
| 14 | Live-Dead | N/A | Sytox Blue | Live-dead discrimination |
| N/A | CD16/32 | 93 | purified | FC receptor blocking antibody |

**Table 2.** Information on Fluorescent Reagents.

| *Parameter* | *Marker* | *Fluorophore* | *Clone* | *Company* | *Catalog Number* | *Stock Concentration* | *RRID* |
| --- | --- | --- | --- | --- | --- | --- | --- |
| 1 | CD45 | BV785 | 30-F11 | BioLegend | 103149 | 0.2 mg/mL | AB_2564590 |
| 2 | CD11b | BV421 | M1/70 | BioLegend | 101235 | 0.05 mg/mL | AB_10897942 |
| 3 | CD11c | PE-Cy7 | N418 | BioLegend | 117317 | 0.2 mg/mL | AB_493569 |
| 4 | F4/80 | PE-Fire 810 | QA17A29 | BioLegend | 157321 | 0.2 mg/mL | AB_3662371 |
| 5 | Ly6G | FITC | 1A8 | BioLegend | 127606 | 0.2 mg/mL | AB_1236494 |
| 6 | Ly6C | BV711 | HK1.4 | BioLegend | 128037 | 0.2 mg/mL | AB_2562630 |
| 7 | SiglecF | PE | S17007L | BioLegend | 155506 | 0.2 mg/mL | AB_2750235 |
| 8 | CD3 | BV750 | 17A2 | BioLegend | 100249 | 0.2 mg/mL | AB_2734148 |
| 9 | CD4 | BV605 | GK1.5 | BioLegend | 100451 | 0.05 mg/mL | AB_2564591 |
| 10 | CD8a | PE-Dazzle-594 | 53-6.7 | BioLegend | 100761 | 0.2 mg/mL | AB_2564026 |
| 11 | CD49b | PerCP-Cy5.5 | HMA2 | BioLegend | 103520 | 0.2 mg/mL | AB_2566105 |
| 12 | $\gamma\delta$TCR | Spark Blue 550 | GL3 | BioLegend | 285241 | 0.2 mg/mL | AB_3662545 |
| 13 | CD19 | BV650 | 1D3/CD19 | BioLegend | 152427 | 0.2 mg/mL | AB_3675002 |
| 14 | CD16/32 | purified | 93 | BioLegend | 101302 | 0.5 mg/mL | AB_312801 |
| 15 | Live-Dead | Sytox Blue | N/A | ThermoFisher | S11348 | 5mM in DMSO | N/A |

**Table 3.** Antibody Cocktail Information.

| *Parameter* | *Marker* | *Stock*<br>*Concentration* | *Volume per*<br>*Sample*<br>*(uL)* | *Amount per*<br>*Sample (ug)* | *Dilution*<br>*During*<br>*Staining* | *Staining*<br>*Concentration*<br>*(ng/uL)* |
| --- | --- | --- | --- | --- | --- | --- |
| 1 | CD45 | 0.2 mg/mL | 2 | 0.4 | 1:125 | 1.6 |
| 2 | CD11b | 0.05 mg/mL | 2.5 | 0.125 | 1:100 | 0.5 |
| 3 | CD11c | 0.2 mg/mL | 2 | 0.4 | 1:125 | 1.6 |
| 4 | F4/80 | 0.2 mg/mL | 3 | 0.6 | ~1:83.33 | 2.4 |
| 5 | Ly6G | 0.5 mg/mL | 1.5 | 0.75 | ~1:166.67 | 3 |
| 6 | Ly6C | 0.2 mg/mL | 2 | 0.4 | 1:125 | 1.6 |
| 7 | SiglecF | 0.2 mg/mL | 2 | 0.4 | 1:125 | 1.6 |
| 8 | CD3 | 0.2 mg/mL | 3 | 0.6 | ~1:83.33 | 2.4 |
| 9 | CD4 | 0.05 mg/mL | 2.5 | 0.125 | 1:100 | 0.5 |
| 10 | CD8a | 0.2 mg/mL | 1.5 | 0.3 | ~1:166.67 | 1.2 |
| 11 | CD49b | 0.2 mg/mL | 3 | 0.6 | ~1:83.33 | 2.4 |
| 12 | gdTCR | 0.2 mg/mL | 3 | 0.6 | ~1:83.33 | 2.4 |
| 13 | CD19 | 0.2 mg/mL | 1.5 | 0.3 | ~1:166.67 | 1.2 |
| 14 | CD16/32 | 0.5 mg/mL | 2.5 | 1.25 | 1:100 | 5 |
| 15 | Live-Dead | 5mM in<br>DMSO | 50uL of 1x solution (1:1000 dilution from stock) |  |  |  |

**Figure 1.**
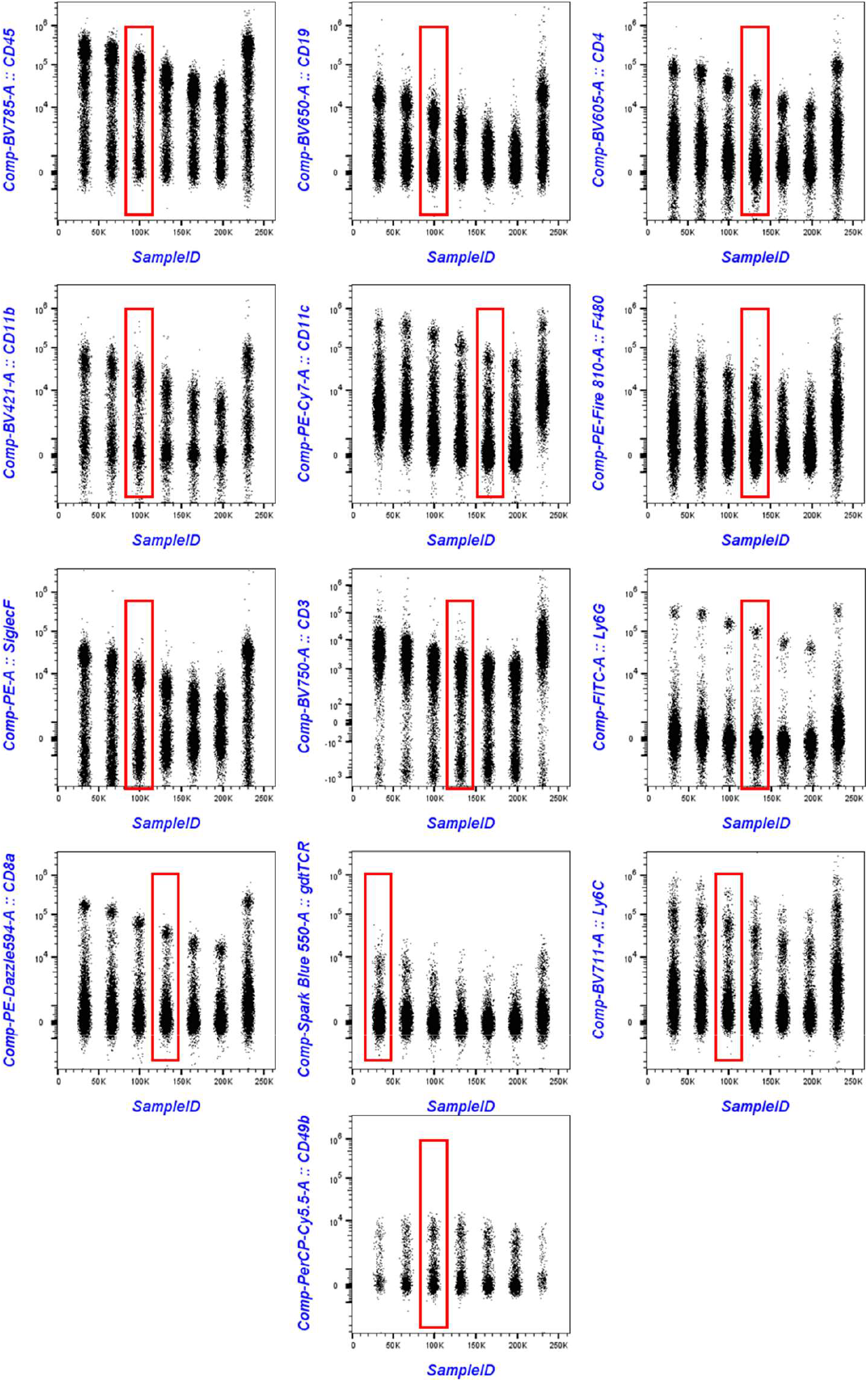
Optimization of antibody dilutions for use in the optimized panel in single cell suspensions from mouse splenocytes. Antibody dilutions are as follows from left to right (1:1, 1:2, 1:4, 1:8, 1:16, 1:32, 2:1) where 1:1 refers to the recommended manufacturer concentration for that antibody. Red box denotes dilution of antibody selected for use in panel.

## Data Acquisition

Samples were analyzed using the 2-laser (Violet, Blue) Cytek Northern Lights Full Spectrum Flow Cytometer. Daily QC was performed using SpectroFlo^®^ QC beads (Lot 2006) prior to acquiring samples to ensure that the cytometer was performing optimally. Daily QC assessed the instrument’s optical alignment and the system performance drift by measuring %rCVs and gains needed to place the beads at the target locations established for each detector. During the typical QC protocol, Laser delays and area scaling factors were also optimized, and gain settings adjusted to account for day-to-day instrument variability. Default QC adjusted instrument settings were used for voltages regarding fluorescent channels, while FSC and SSC parameters were set as follows based on our experience with mouse BALF cells (**FSC:** 50, **SSC:** 112, **SSC B:** 97). These settings may be instrument specific and it is recommended that investigators utilize their knowledge of leukocyte characteristics to adjust these parameters to obtain the best look at the leukocyte populations. See **Table 4** for instrument configuration and peak fluorophore channels. Spectral unmixing with autofluorescence extraction was performed using the SpectroFlo version 3.03 software, with single stained reference controls for each fluorophore as well as an unstained sample. All reference controls were treated the same as the BALF samples. Data were then analyzed using FlowJo Software version 10.10.0. For dimensionality reduction analysis, files of representative animals for each timepoint were pre-gated to remove debris, doublets and dead cells and select only CD45+ leukocyte populations prior to concatenation with preservation of SampleID and SpecimenID using the built in function in FlowJo version 10.10.0. Dimensionality reduction was performed using default tSNE settings on the built in opt-SNE algorithm^17^ in FlowJo version 10.10.0 using 1000 iterations, perplexity of 30, automatically calculated learning rate for the sample. KNN algorithm: Exact (vantage point tree) and gradient algorithm: FFT Interpolation (Fit-SNE)^18^.

**Table 4.** Peak Fluorophore Channels for Reagents Used in this Panel.

| Laser | Channel | Center Wavelength | Bandwidth (nm) | Wavelength Start (nm) | Wavelength End (nm) | Fluorophore Peak |
| --- | --- | --- | --- | --- | --- | --- |
| Violet Laser | V1 | 428 | 15 | 420 | 435 | BV421 |
|  | V2 | 443 | 15 | 436 | 451 |  |
|  | V3 | 458 | 15 | 451 | 466 |  |
|  | V4 | 473 | 15 | 466 | 481 | Sytox-Blue |
|  | V5 | 508 | 20 | 498 | 518 |  |
|  | V6 | 525 | 17 | 516 | 533 |  |
|  | V7 | 542 | 17 | 533 | 550 |  |
|  | V8 | 581 | 19 | 571 | 590 |  |
|  | V9 | 598 | 20 | 588 | 608 |  |
|  | V10 | 615 | 20 | 605 | 625 | BV605 |
|  | V11 | 664 | 27 | 651 | 678 | BV650 |
|  | V12 | 692 | 28 | 678 | 706 |  |
|  | V13 | 720 | 29 | 706 | 735 | BV711 |
|  | V14 | 750 | 30 | 735 | 765 | BV750 |
|  | V15 | 780 | 30 | 765 | 795 | BV785 |
|  | V16 | 812 | 34 | 795 | 829 |  |
| Blue Laser | B1 | 508 | 20 | 498 | 518 |  |
|  | B2 | 525 | 17 | 516 | 533 | FITC |
|  | B3 | 542 | 17 | 533 | 550 | Spark Blue 550 |
|  | B4 | 581 | 19 | 571 | 590 | PE |
|  | B5 | 598 | 20 | 588 | 608 |  |
|  | B6 | 615 | 20 | 605 | 625 | PE-Dazzle 594 |
|  | B7 | 661 | 17 | 653 | 670 |  |
|  | B8 | 679 | 18 | 670 | 688 |  |
|  | B9 | 697 | 19 | 688 | 707 | PerCP-Cy5.5 |
|  | B10 | 717 | 20 | 707 | 727 |  |
|  | B11 | 738 | 21 | 728 | 749 |  |
|  | B12 | 760 | 23 | 749 | 772 |  |
|  | B13 | 783 | 23 | 772 | 795 | PE-Cy7 |
|  | B14 | 812 | 34 | 795 | 829 | PE-Fire 810 |

## Panel Information, Gating Strategy and Population Identification

The reagents used for our panel are described in **Table 1**. These fluorophores are excitable by the Violet or Blue Lasers and allow for use of this panel on a 2-laser full spectrum cytometer such as the Cytek Northern Lights VB2000 cytometer we utilized here. This panel can be easily adapted for other full spectrum cytometers provided they have blue and violet lasers. Analysis begins with pre-gating by excluding debris, doublets, dead cells, and non-leukocytes as shown in **Figure 2A**. Briefly, debris is excluded by gating cells based on FSC-A and SSC-A parameters. From this gate, we narrow in on Live Leukocytes by selecting CD45^+^ Sytox Blue^-^ (viability dye) cells. Doublets are removed using FSC-A vs FSC-H, making sure to be flexible with gating on the larger cells (macrophages) for which the area does not scale perfectly linear with height. After gating on live single leukocytes, we then begin by identifying neutrophil populations by plotting the Live Single Leukocytes on CD11c vs Ly6G and identifying neutrophils as CD11c^-/lo^ and Ly6G^+^. The remainder of the population is then gated on CD11c vs SiglecF, where eosinophils are identified as CD11c^-^ SiglecF^+^ cells that are further confirmed to be FSC-A^lo^ and SSC-A^hi^ as small but highly granular cells. Following removal of neutrophils and eosinophils from Live Single Leukocytes, the remaining cells are plotted as CD11b vs F4/80 to distinguish lymphoid cells from monocytes and macrophages. The lymphoid gate is further confirmed to be FSC-A^lo^ SSC-A^lo^ cells characteristic of lymphocytes, then is plotted as CD19 vs gamma-delta (γδ) TCR to identify B cells (CD19^+^) and γδT cells (γδ TCR^+^). The rest of the lymphoid cells are then plotted as CD3 vs CD49b to identify NK cells (CD3^-^CD49b^+^) and T and NKT cells (CD3^+^CD49b^+/-^) which can be further subdivided into helper T and cytotoxic T cells based on CD4 and CD8 expression profiles **Figure 2B**. Tissue resident alveolar macrophages (resMacs) versus monocyte derived (recMacs) can be further distinguished based on SiglecF, where tissue resident macrophages are SiglecF^hi^ whereas monocyte derived macrophages are SiglecF^lo/-,^ based on previously published strategies^19^ **Figure 2C**. More subpopulations can be identified by further gating, however these subpopulations cannot be confirmed with 100% certainty. For example, recMacs can be further subdivided into CD11b+ Ly6C+ which can represent newly recruited inflammatory (classical) monocytes and CD11b+Ly6C-populations which can represent either non-classical monocytes or classical transitioning monocytes that begin to replenish the lost tissue resident alveolar macrophage niche. resMacs can be subdivided into CD11b+ activated tissue resident alveolar macrophages or CD11b-resting tissue resident alveolar macrophages. Furthermore, CD49b can be utilized to further subdivide the CD3+ T cell populations into NKT like cells, however as CD49b can be upregulated in conventional T cells during inflammation it cannot be stated with full confidence whether CD49b+CD3+ T cells are NKT cells or simply conventional T cell subsets that are activated. Major markers used to confidently identify major populations are described in the table below. More subsets can be identified then shown using this panel, but not with a high degree of certainty (ex. Dendritic cells (DCs) as F4/80^-^, CD11c^+^, CD11b^+/-^, Classical Monocytes CD45^+^ CD11c^-^CD11b^+^ Ly6C^+^ SiglecF^-^, NK cells can be classified as mature or immature based on Ly6C expression, etc.). Marker combinations used for the identification of primary cell populations are outlined in **Table 5** below. Dimensionality reduction via opt-SNE^17,18^ can be applied to enable 2-dimensional comparisons between different samples, and we used it here to identify specific diferences in host responses in mice intranasally infected with three different species of Mycoplasmas compared to uninfected mice, as well as to phenotype the evolution of the inflammatory landscape in the airway over a 21 day time course in infected mice (**Figure 3** and **4**).

**Table 5.**
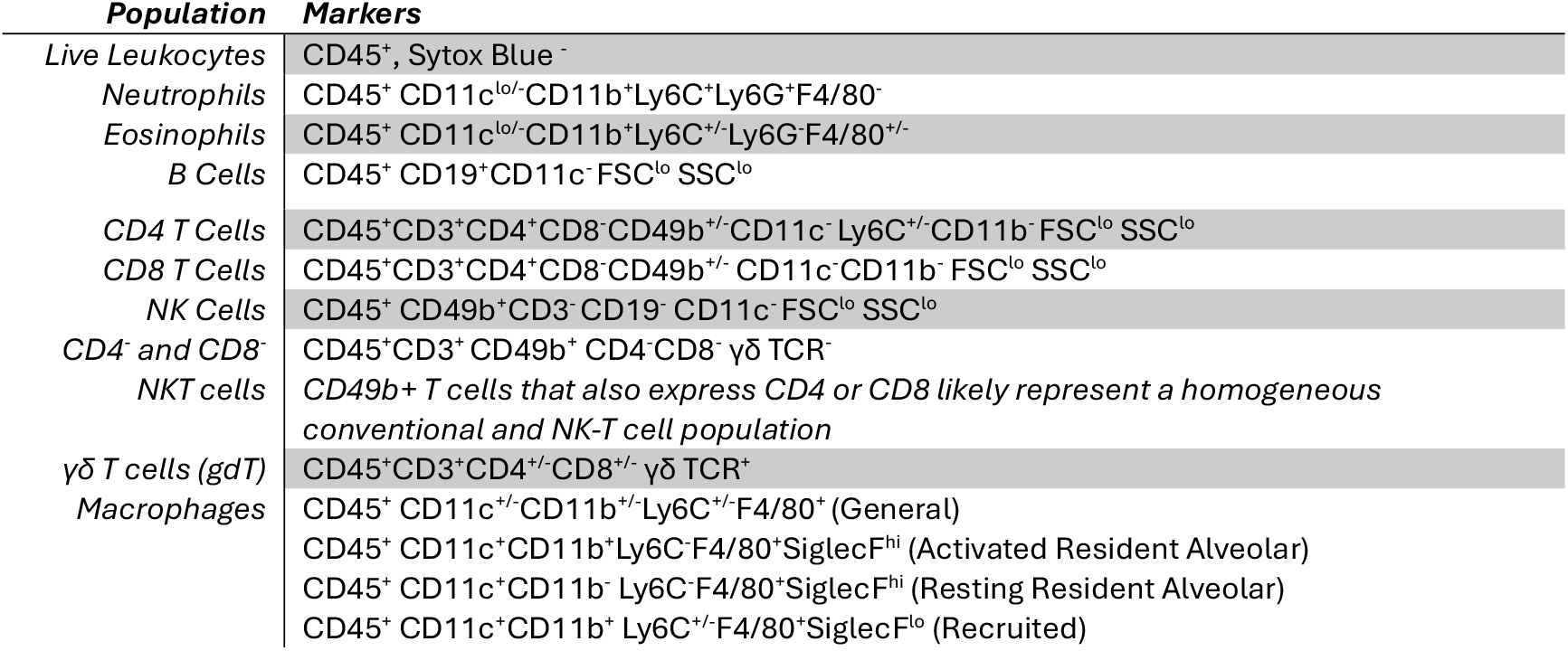
Marker combinations for identification of major leukocytes.

**Figure 2.**
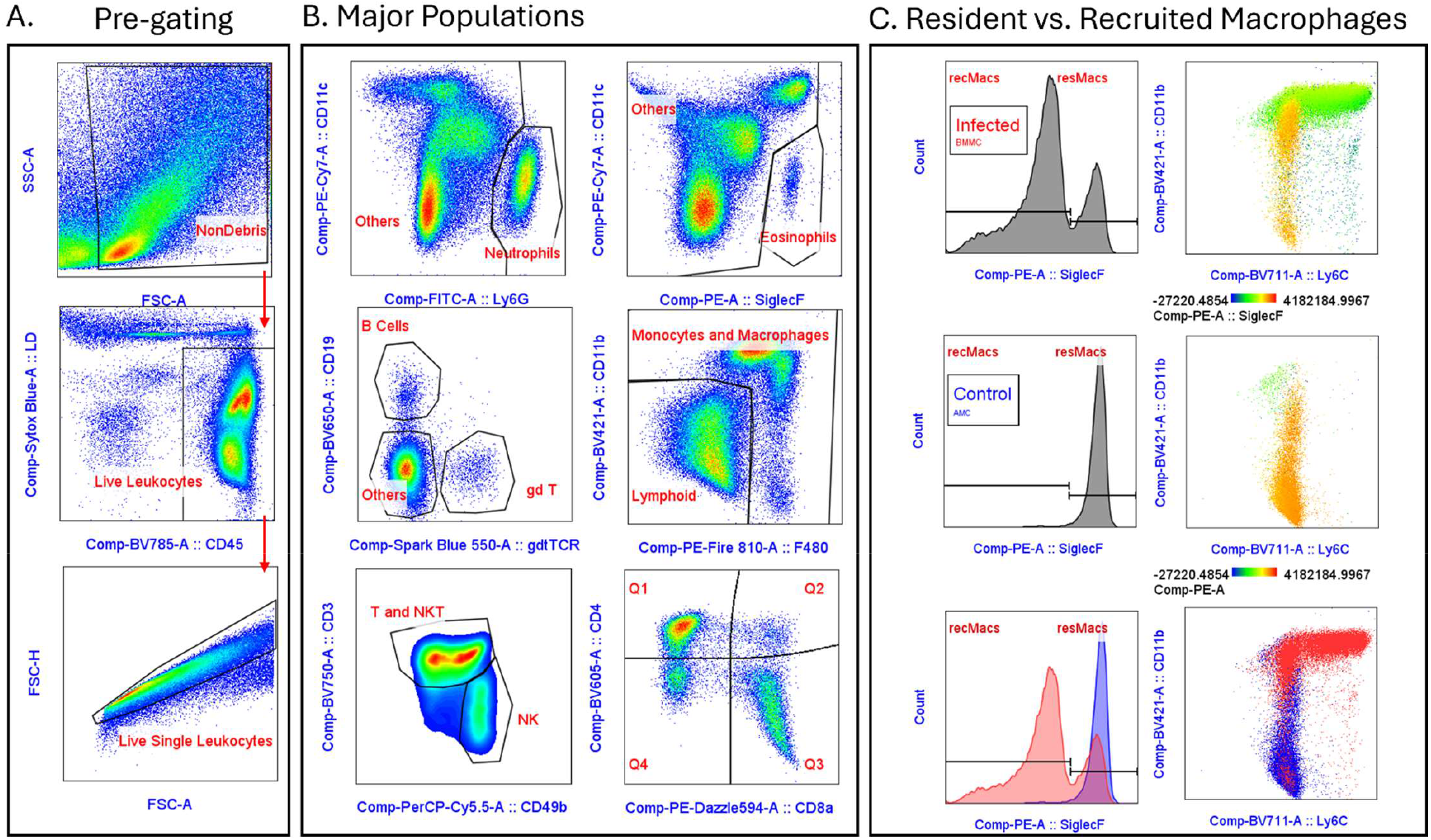
Representative gating strategy used to identify major cell populations in the BALFs of infected mice. **(A)** Pre-gating strategy that removes debris, doublets and homes in on Live Single Leukocytes for further downstream analysis. **(B)** Gating of major cell populations. **(C)** Distinguishing between tissue resident versus recruited monocyte derived macrophages using SiglecF expression profiles. Note that conventional gating can reveal further subsets of cell populations that are not shown in the gating strategy above.

**Figure 3.**
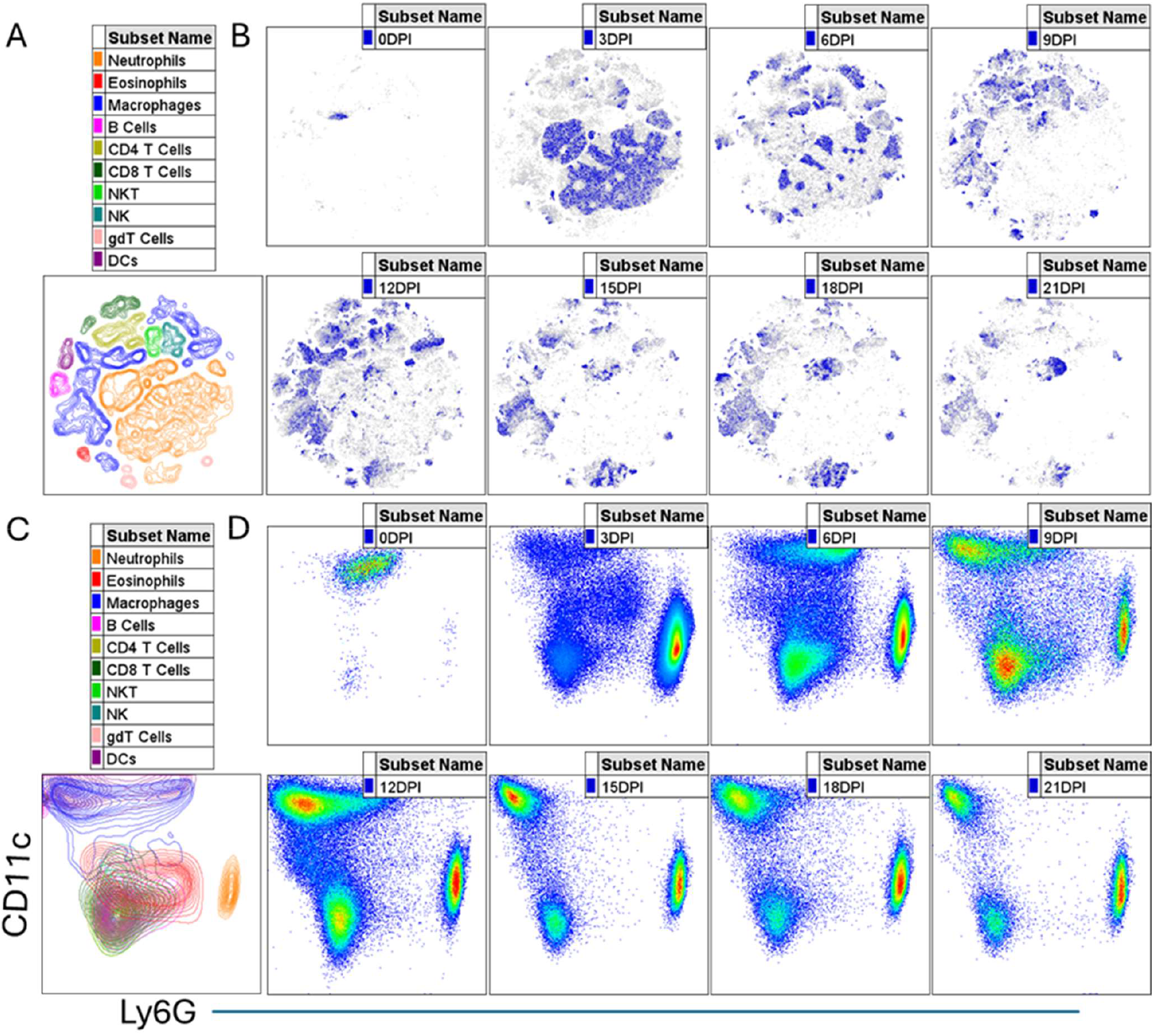
Longitudinal assessment of airway inflammation in mice infected with Mycoplasma pneumoniae using our panel. **(A)** Identified cell clusters following dimensionality reduction via opt-SNE (see methods). **(B)** Cell density plots of representative samples per timepoint indicating the phenotype of airway inflammation. **(C)** Identified cell clusters as in A but represented on a CD11c vs Ly6G two-dimensional plot. **(D)** Pseudocolor density plot of representative sample per timepoint demonstrating phenotype of airway inflammation on two-dimensional CD11c vs. Ly6G plots.

**Figure 4.**
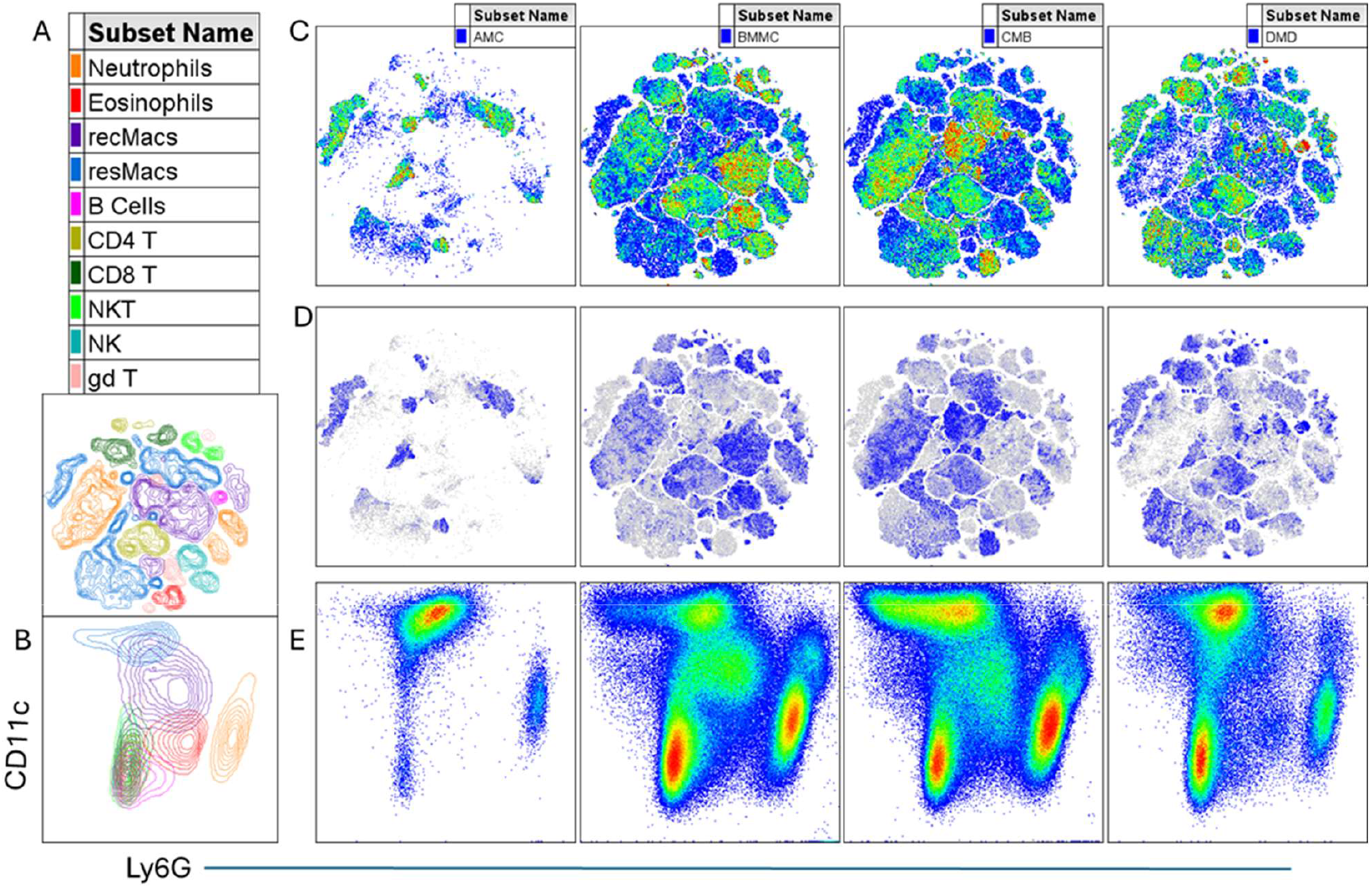
Comparative Immunophenotyping of airway inflammation induced by intranasal infection with: AMC (sterile growth medium), BMMC (Mycoplasma mycoides subsp. capri), CMB (Mycoplasma bovis), DMD (Mycoplasma dispar). **(A)** Identified cell clusters following dimensionality reduction via opt-SNE (see methods). **(B)** Identified cell clusters as in A but represented on a CD11c vs Ly6G two-dimensional plot. **(C)** Pseudocolor and **(D)** cell density plots of representative samples of mice infected with different Mycoplasma spp. **(E)** Pseudocolor density plot of representative samples demonstrating phenotype of airway inflammation on two-dimensional CD11c vs. Ly6G plots.

Cell clusters can be identified with high confidence and verified using conventional gating strategies.

## Important Panel Notes

F4/80 clone BM8 did not provide good separation of macrophage populations and was first replaced with anti-mouse CD64 clone S18017D on PE/Fire – 744. This provided good macrophage separation however the fluorophore was prone to conjugate breakdown if MasterMix was not prepared on the same day, whereas anti-F4/80 antibody clone QA17A29 on PE-Fire 810 provided good separation of macrophage populations enabling better Dendritic cell identification than clone BM8 and the reagent was found to be more stable in stored MasterMix for up to 1 week. Addition of anti-MHC-II on BV510 did provide better resolution for more confident identification of dendritic cells but introduced unmixing artefacts that required more advanced unmixing methods than what is automatically provided by the software. As our goal was to present a panel that can be easily adopted with minimal end-user optimization, we chose to move forward with the current version of the panel that we describe in this manuscript. Clone 6D5 of anti-mouse CD19 antibody did not provide good separation for our experiments and we recommend that clone 1D3/CD19 is used instead, as described in our panel. The evolution of this panel is detailed in **Table 6** below.

**Table 6.** Panel Evolution and Iterations.

| **Parameters** | **Version 1** | **Version 2** | **Version 3** | **Version 4** | **Version 5** | **Current** |
| --- | --- | --- | --- | --- | --- | --- |
| 1 | CD45 | CD45 | CD45 | CD45 | CD45 | CD45 |
| 2 | CD11b | CD11b | CD11b | CD11b | CD11b | CD11b |
| 3 | CD11c | CD11c | CD11c | CD11c | CD11c | CD11c |
| 4 |  | F4/80<br>(clone<br>BM8) | CD64<br>(clone<br>S18017D:<br>PE/Fire<br>744) | CD64<br>(clone<br>S18017D:<br>PE/Fire 744) | CD64<br>(clone<br>S18017D:<br>PE/Fire 744) | F4/80<br>(current<br>clone<br>described<br>above) |
| 5 | Ly6G | Ly6G | Ly6G | Ly6G | Ly6G | Ly6G |
| 6 | Ly6C | Ly6C | Ly6C | Ly6C | Ly6C | Ly6C |
| 7 | SiglecF | SiglecF | SiglecF | SiglecF | SiglecF | SiglecF |
| 8 |  | CD3 | CD3 | CD3 | CD3 | CD3 |
| 9 | CD4 | CD4 | CD4 | CD4 | CD4 | CD4 |
| 10 | CD8a | CD8a | CD8a | CD8a | CD8a | CD8a |
| 11 |  |  | CD49b | CD49b | CD49b | CD49b |
| 12 |  |  |  |  | gdTCR | gdTCR |
| 13 | CD19 | CD19 | CD19 | CD19 | CD19 | CD19 |
| 14 | CD16/32 | CD16/32 | CD16/32 | CD16/32 | CD16/32 | CD16/32 |
| 15 | Live Dead | Live Dead | Live Dead | Live Dead | Live Dead | Live Dead |
| 16 |  |  |  | MHC-II /I-A/I-E (clone<br>M5/114.15.2,<br>BV510) | MHC-II /I-A/I-E (clone<br>M5/114.15.2,<br>BV510) |  |

## Conclusion

Our optimized 14-color BALF immunophenotyping panel offers a practical, accessible and biologically comprehensive solution for studying airway inflammation in murine models. By balancing depth of resolution with ease of use, it enables consistent identification of key myeloid and lymphoid populations across diverse experimental conditions, supports both discovery-driven and mechanistic studies, and integrates seamlessly with downstream functional or transcriptomic workflows. Its demonstrated utility across multiple respiratory Mycoplasma infection models underscores its robustness and broad applicability. Together, these features position the panel as a valuable community resource that can standardize analyses, enhance reproducibility, and accellerate comparative insights into pulmonary immunity.

## Author Contributions

### Arlind B. Mara

Conceptualization, data curation, formal analysis, experimentation, investigation, methodology, validation, visualization, writing (original draft, review & editing). **Jeremy M. Miller:** Experimentation, investigation, writing (review & editing). **R. Grace Ozyck and Morgan L Hunte**: Experimentation and animal sampling. **Edan R. Tulman, Steven M. Szczepanek and Steven J. Geary:** Supervision, funding acquisition, resources, writing (review & editing).

## Acknowledgments

The work described herein was supported by funds made available by the Center of Excellence for Vaccine Research (SJG) and the USDA-ARS Non-Cooperative Agreement #5030-32000-236-002S.

## Data Availability Statement

The datasets associated with this manuscript have been made publicly available in the *Zenodo* repository and can be accessed through the following link: https://doi.org/10.5281/zenodo.17514115.

